# Experimental evolution of a pheromone signal

**DOI:** 10.1101/2021.09.06.459111

**Authors:** Thomas Blankers, Elise Fruitet, Emily Burdfield-Steel, Astrid T. Groot

## Abstract

Sexual signals are important in speciation, but understanding their evolution is complex as these signals are often composed of multiple, genetically interdependent components. To understand how signals evolve, we thus need to consider selection responses in multiple components and account for the genetic correlations among components. One intriguing possibility is that selection changes the genetic covariance structure of a multicomponent signal in a way that facilitates a response to selection. However, this hypothesis remains largely untested empirically. In this study, we investigate the evolutionary response of the multicomponent female sex pheromone blend of the moth *Heliothis subflexa* to 10 generations of artificial selection. We observed a selection response of about 3/4s of a phenotypic standard deviation in the components under selection. Interestingly, other pheromone components that are biochemically and genetically linked to the components under selection did not change. We also found that after the onset of selection, the genetic covariance structure diverged, resulting in the disassociation of components under selection and components not under selection across the first two genetic principle components. Our findings provide rare empirical support for an intriguing mechanism by which a sexual signal can respond to selection without possible constraints from indirect selection responses.

## INTRODUCTION

Animal taxa that engage in sexual communication typically show high among-species diversity in sexual signals (Andersson, 1994; Coyne and Orr, 2004; Ritchie, 2007; Schaefer and Ruxton, 2015; Wiens and Tuschhoff, 2020). This diversity in sexual signals is generally hypothesized to be due to directional or disruptive selection (Ritchie, 2007; Schaefer and Ruxton, 2015; West-Eberhard, 2014; Wilkins et al., 2013), although sexual signal evolution is still a mystery in many species. Since sexual signals play an important role in the origin and maintenance of species and contribute to biodiversity (Coyne and Orr, 2004), it is important to assess whether there are constraints to their selection response and identify the mechanisms that can mitigate those constraints.

Understanding selection responses in sexual signals is challenging, because signals are often composed of multiple components (Candolin, 2003; Higham and Hebets, 2013; Rowe, 1999). For example, mating songs can vary both in pitch and in rhythm (Blankers et al., 2015; Tanner et al., 2017; Wilkins et al., 2015), color signals can be composed of multiple, functionally distinct patches (Cole and Endler, 2015; Grether et al., 2004), and sex pheromones are often blends of multiple chemical compounds (Ferveur, 2005; Linn et al., 1987). To understand how multicomponent signals evolve, we thus need to consider the selection response in multiple dimensions simultaneously. Moreover, signal components can have a shared genetic or developmental basis, or can be subject to correlated selection pressures (Armbruster et al., 2014; Cheverud, 1996). The resulting genetic correlations among signal components can influence how selection on the phenotype translates to changes in the underlying genotypes (Chenoweth and Blows, 2006). To understand how the genotype-phenotype map of sexual signals influences the selection response, we thus need to determine the genetic correlations between the different signal components.

Statistical frameworks in quantitative genetics, in particular the (multivariate) breeder’s equation, allow us to predict and quantitatively understand selection responses in correlated traits. In this framework, the response to selection is a function of the genetic (G) and phenotypic (P) variance-covariance matrix and the selection gradient: selection acts on the P matrix and the resulting response is constrained by the G matrix (Lande, 1979; Lande and Arnold, 1983; Lynch and Walsh, 1998). The difficulty in predicting selection responses of multivariate traits is that selection acting on multiple components may be counterbalancing, e.g. directional selection on one trait, but stabilizing selection on correlated traits, resulting in evolutionary constraints (Barton and Turelli, 1989). Counterbalancing selection is likely prevalent in the evolution of sexual signals, as choosing individuals may favor higher or lower values of some component of the signal, while changes in correlated components may result in reduced mate attraction.

These evolutionary constraints can be overcome if genetic variances and covariances themselves respond to selection, thus reshaping the G matrix (Arnold et al., 2008; Barton and Turelli, 1989; Eroukhmanoff, 2009; Jones et al., 2003; Melo and Marroig, 2014; Revell, 2007; Roff and Fairbairn, 2012). Theory predicts that the G matrix will vary through time, because on one hand selection erodes genetic variance (Barton and Turelli, 1989; Estes and Arnold, 2007), while on the other hand mutation and introgression add new variation, albeit more slowly. Moreover, genetic correlations can respond to selection directly, especially if they result from interactions among unlinked genetic loci affecting the co-expression of multiple traits (Wolf et al., 2005), or from selection acting on correlations directly (Armbruster et al., 2014; Roff and Fairbairn, 2012; Svensson et al., 2021). Interestingly, directional selection can change genetic covariances and increase modularity, meaning that groups (modules) of traits become genetically independent from other modules, in a way that facilitates the phenotypic response to selection (Melo and Marroig, 2014). Empirical work has shown that patterns of covariation can indeed evolve both across populations and time in nature (Bégin and Roff, 2003; Berner et al., 2010; Björklund et al., 2013; Blankers et al., 2017; Garant et al., 2008; Gosden and Chenoweth, 2014) and during artificial selection experiments (Careau et al., 2015; Hine et al., 2011; Uesugi et al., 2017). However, it is still unclear how changes in the phenotypic selection response are related to changes in genetic (co)variance through time.

In this study, we explored the response in phenotypic means and genetic (co)variances to artificial selection on the female sex pheromone of the moth *Heliothis subflexa* (Lepidoptera, Noctuidae). Like many other moths, *H. subflexa* females secrete a sex pheromone blend to which conspecific males are attracted. These sex pheromone blends are species-specific and vary among species in both the presence/absence of components as well as in relative amounts (or ratios) of the components (Cardé and Haynes, 2004; Schneider, 1992).

The sex pheromone blend of *H. subflexa* females consists of 11 compounds, with the following components that are critically important for conspecific male attraction: (Z)-11-hexadecenal (Z11-16:Ald) as the major sex pheromone component, (Z)-9-hexadecenal (Z9-16:Ald) and (Z)-11-hexadecenol (Z11-16:OH) as the two secondary sex pheromone components, without which *H. subflexa* males are not attracted (Groot et al., 2007; Vickers, 2002). Interestingly, the acetate esters (Z)-7-hexadecenyl acetate (Z7-16:OAc), (Z)-9-hexadecenyl acetate (Z9-16:OAc), and (Z)-11-hexadecenyl acetate (Z11-16:OAc), from here on referred to as “acetates”, have a dual role: these acetates attract conspecific males, while simultaneously repelling males of *H. virescens* (Groot et al., 2006; Vickers and Baker, 1997). In geographic regions where *H. virescens* is present, acetates are more abundant in the *H. subflexa* pheromone compared to where this species is absent (Groot et al., 2009a). This suggests that the acetates are subject to divergent selection across a geographic cline. The relative amounts of the other components are hypothesized to be under stabilizing selection across the range, as in general in moth pheromone communication the mean blend is preferred over deviations from the mean (Allison and Cardé, 2008; Groot et al., 2010; Kárpáti et al., 2013; Linn et al., 1997; Löfstedt, 1990; Zhu et al., 1997).

To explore the selection response of the acetates, we selected for higher and lower amounts of acetates during 10 generations of truncation selection. Since the acetates vary geographically (Groot et al., 2009a) and have a genetic basis that is partially independent of other components (Groot et al., 2009b), we hypothesized that the relative amount of acetates can evolve in response to univariate selection for higher/lower acetates but that genetic variance will be reduced after selection. However, since acetates also partially share their genetic basis with other components and since all sex pheromone compounds are produced through the same biosynthetic pathway, we also hypothesized that there will be indirect selection responses in other pheromone components, specifically in the unsaturated aldehydes, Z9-16:Ald and Z11-16:Ald, and alcohol, Z11-16:OH (Fig 1). Since some of the correlations among components may reduce the selection response, we also expected genetic covariances to change during selection, specifically in a way that facilitates the phenotypic selection response.

**Figure 1.**
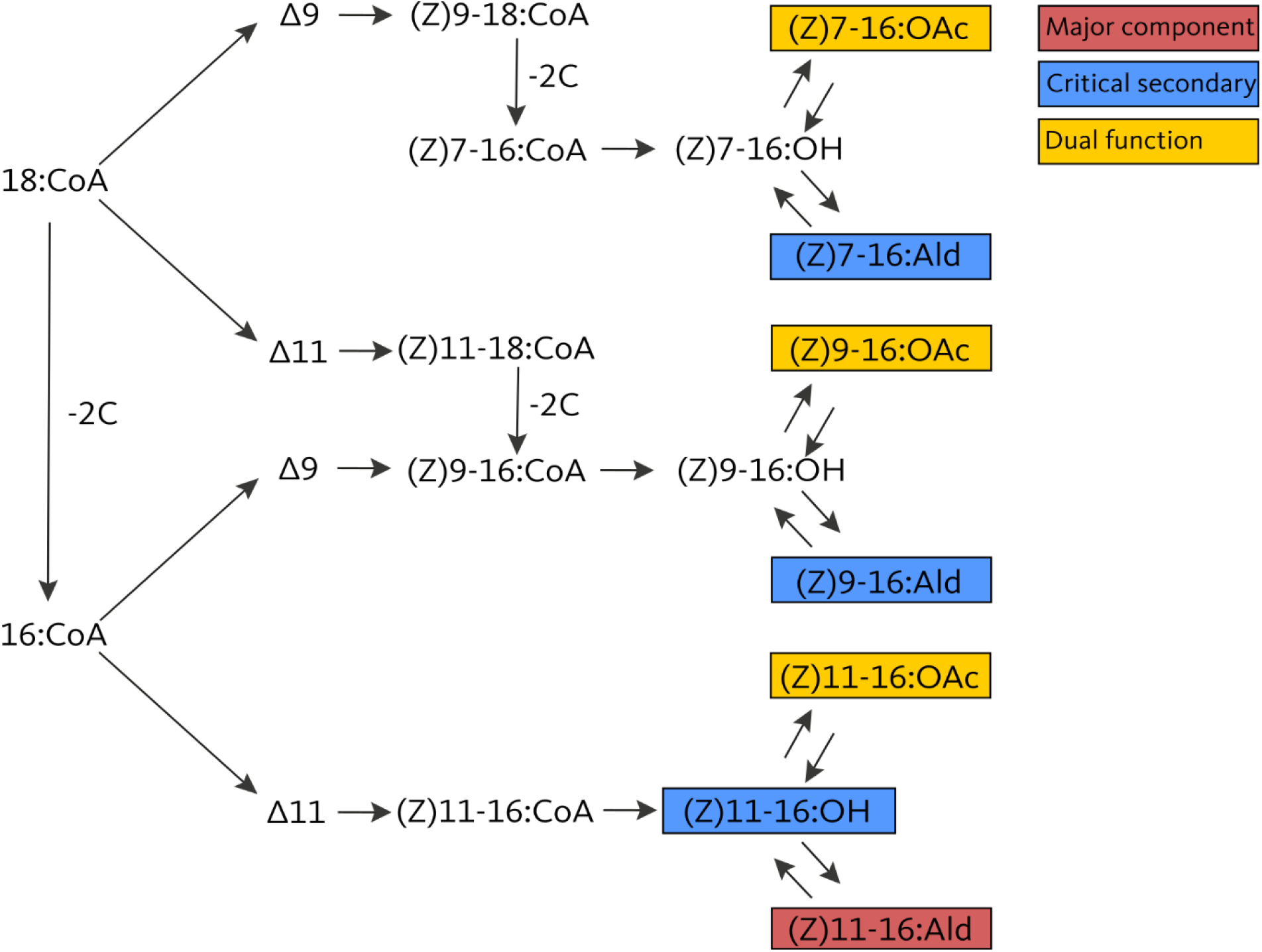
Simplified biosynthetic pathway of the *Heliothis subflexa* sex pheromone, redrawn from (Groot et al., 2009b) and based on (Jurenka, 2003). Desaturation and β-oxidation produce mono-unsaturated acyl-CoA precursors from 18 or 16 carbon acyl CoA derivatives, which are then modified to form acetates, alcohols, and aldehydes through specific enzymatic conversions. The Δ9 and Δ11 desaturases create a double bond between the 9^th^ and 10^th^ or 11^th^ and 12^th^ carbon of the 18-or 16-carbon acyl-CoA derivatives, respectively. β-oxidation shortens the chain length from 18 to 16 carbons. Note that (Z)9-16 acyl-CoA can be formed through two alternative routes. The compounds present in the pheromone glands of *H. subflexa* are in boxes that are color coded depending on their function in male response behavior.

## METHODS

### Insect rearing

The laboratory population of *H. subflexa* was founded from animals collected in the field in 2006 and has been reared at the University of Amsterdam since 2011 using single pair matings to maintain genetic diversity. There has been occasional exchange among *H. subflexa* populations at North Carolina State University, Amsterdam, and the Max Planck Institute for Chemical Ecology in Jena. Eggs collected from single pair matings were kept in Petri dishes (Ø 85 mm) with artificial wheat germ/soy flour-based diet (BioServ Inc., Newark, DE, USA) at room temperature for approximately 10 days, after which larvae were reared in separate individual 37-mL cups filled with the same artificial diet and kept at 25°C and 60% relative humidity with 14h:10h light-dark cycle. Upon emergence, adults were provided sugar water. Pairs of males and females were housed in 375 mL paper cups covered with gauze and kept under the same conditions as the late-instar larvae and virgin adults. The mating pairs were provided with sugar water. To stimulate oviposition, a freshly cut gooseberry fruit was placed on top of the gauze. Once the eggs began to hatch, the gauze and the eggs and larvae on it, were transferred to Petri dishes, which were then placed at room temperature until larvae were transferred to individual cups. The females that produced fertile offspring were collected and phenotyped, if still alive. All matings were assigned unique numbers. This way, we obtained a full pedigree of all individuals in the selection and control lines.

### Phenotyping

Female sex pheromone can be extracted from mated and old females by injecting them with Pheromone Biosynthesis Activating Neuropeptide, i.e. PBAN (Groot et al., 2005). Females were injected with a 7.5 pmol (2 µL of a 0.0146 µg/µL) PBAN solution to activate pheromone production post mating. After a 90-minute incubation time, female glands were extruded by squeezing the abdomen then fixed by firmly holding the abdomen with forceps just anterior of the gland. The gland was excised with microdissection scissors, and the moths were euthanized. Excess abdominal tissue and eggs that remained in the ovipositor were removed, after which the glands were submerged in 50 μL hexane containing 200 ng pentadecane as an internal standard. After 30 minutes, the glands were removed and the extracts stored at −20°C until analysis.

Pheromone extracts were analyzed by injecting the concentrated samples into a splitless inlet of a 7890A GC (Agilent Technologies, Santa Clara, CA, USA). The area under the pheromone peaks was calculated using integration software implemented in Agilent ChemStation (version B.04.03). Pheromone peak areas were obtained for the 11 pheromone components: two 14-C aldehydes (14:Ald, Z7-14:Ald), four 16-C aldehydes (16:Ald, Z7-16:Ald, Z9-16:Ald, Z11-16:Ald), the three acetates (Z7-16:OAc, Z9-16:OAc, Z11-16:OAc), and two 16-C alcohols (Z9-16:OH, Z11-16:OH). Z7-16:Ald and Z9-16:Ald were difficult to separate by GC and were therefore integrated as one peak (referred to Z9-16:Ald). Absolute amounts (in ng) of each compound were calculated relative to a 200 ng pentadecane internal standard. All downstream analyses were done in R 3.6.1 (R Core Team, 2019). Samples containing < 20 ng were excluded, because the ratios of the components in such low titers cannot be reliably measured in the chromatogram. Relative amounts were calculated by dividing the absolute amounts by the total amount across all 11 components.

### Data transformation

Selection was performed based on relative amounts of acetates. However, describing relationships among relative amounts is problematic, because they sum to 100%, thereby mathematically constraining the (co)variation in the pheromone and biasing the analysis. We therefore transformed pheromone measurements to log-contrasts for all down-stream analyses in this study. This approach breaks the interdependency and normalizes the data. Since the divisor used in the contrast of the variable can no longer be part of any downstream analyses, we chose 14:Ald as the divisor, because this component has a small but clearly detectable peak in the chromatogram, while it is irrelevant for male response behavior (Heath et al., 1990). Prior to downstream analyses, samples with a χ^2^-distributed Mahalanobis distance score (calculated using the ‘mahalanobis’ function in the ‘stats’ package) that exceeded a threshold value corresponding to a Bonferroni-corrected P-value < 0.05 were removed, which resulted in removing 64 out of a total of 2,861 samples. These samples showed abnormal pheromone ratios and were present across all selection and control lines and generations. We are therefore confident that these samples represent outliers, e.g. due to extraction or measurement errors, and are not representative of any relevant biological phenomena.

### Selection

Since phenotyping consisted of extracting the sex pheromone gland invasively, selection was performed post-mating. Each generation, we continued with those families that had a maternal phenotype satisfying an increasingly stringent threshold. Initially, a so-called “high” and “low” line were started with offspring from females with a relative amount of acetates above 22% or below 16%, respectively. These values were based on the distribution of the relative amounts of acetates in the starting population, representing the 1st and 3rd quantile. These thresholds were kept for the first 3 generations. In subsequent generations, we increased these thresholds to >24% or <14% (generation 4 – 5), and >26% or <12% (generation 6 – 9). The high line consisted of a total of 2,236 breeding females across nine generations, ranging from 189 – 295 matings per generation. After outlier removal, a total of 1,234 high line females were phenotyped, or between 89 and 171 per generation during the selection phase. The low line consisted of 2,250 breeding females (189 – 296 per generation) and, after outlier removal, phenotypes were measured for a total of 1,180 females (57 – 169 per generation). Because maintenance of the lines required individual rearing (due to larval cannibalism) and phenotyping of so many individuals, we were unable to maintain replicate lines of the high and low line. Parallel results between the lines in the selection response thus lend confidence to the patterns observed. However, we avoid drawing conclusions based on differences between the high and low lines, because these can both reflect differential selection effects or sampling variance. We stopped selecting in generation 10, but continued to phenotype ∼25 females per line and per generation until generation 13. Throughout the selection experiment, we also maintained a control line from which we phenotyped between 6 and 111 females (median 20 females) per generation. For detailed sample sizes see Table A1 in the appendix.

### Phenotypic selection response

To test the hypothesis that acetates can respond to selection and that other pheromone components change due to indirect selection, the selection responses as measured by the log-contrasts were visualized using the R-package ‘ggplot’. In addition, we also calculated the average difference between selected and control lines in units of standard deviation of the starting population. To test whether log-contrast ratios for pheromone components had significantly increased or decreased during selection, we compared the starting generation with the final generation for the control, high, and low line using a Student’s t-test.

### Genetic (co)variance selection response

To test whether genetic variances decreased with selection and whether the structure of the genetic variance-covariance matrix was affected by selection, we first ran multi-response animal models implemented in the R-package ‘MCMCglmm’. We limited our analyses to the four components that have been shown to affect male response: Z9-16:Ald, Z11-16:Ald, Z11-16:OAc, and Z11-16:OH (in all cases, the log-contrast to 14:Ald was used). Model formulation roughly followed the MCMCglmm manual (Hadfield, 2010) and Jarrod Hadfield’s course notes (Hadfield, 2012), as well as the animal model tutorial by Pierre de Villemereuil (Villemereuil, 2012). High and low line individuals had separate pedigrees, but shared ancestors for which we had pedigree data up to three generations prior to the onset of the experiment (in total 1,065 breeding females). To minimize the influence from priors on the covariances, for which we had no prior expectations with high degrees of belief, we formulated flat, uninformative priors for the variance-covariance structure of the random effect (‘animal’). To evaluate the models, we checked chain convergence and assessed effective sample sizes and levels of autocorrelation. To check whether the prior did not contribute unduly to posterior estimates, we also compared genetic (co)variance estimates among univariate, bi-variate, and full (tetra-variate) models and using different, more informative priors. We found models with flat priors to perform best.

We obtained estimates of genetic (co)variances for the starting populations (generation zero and one combined), for generation two and three combined (early response) and for generation nine (final response). For each generation, or pair of generations, we obtained posterior estimates of the additive genetic variance, V_A_ for each of the four log-contrasts from the multi-response animal model. For each posterior sample of V_A_, we calculated the coefficient of additive genetic variance, 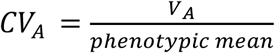, which provides a standardized measure of the evolvability of a trait that is independent of other variance components. We also examined changes in the G matrix by inspecting the trait loadings on genetic principle components. The four genetic principle components were obtained for every posterior sample using the ‘eigen’ function. We focused on the first two axes, gPC1 (also known as *g*_*max*_, the direction in phenotypic space containing the largest fraction of the genetic variance) and gPC2, because these axes jointly described > 90% of the genetic variance (see Results). Comparisons of posterior distributions was done using the Honest Posterior Density. Posteriors were considered statistically significantly different if 90% HPD intervals did not overlap.

## RESULTS

### Selection response in sex pheromone

All three acetates responded to selection for higher/lower relative total amounts of acetates in the high and low line, respectively (Fig 2). The line-specific means significantly increased in the high line for all three acetates between generation 0 and generation 9, and significantly decreased in the low line, while no significant differences were observed for the back-up line (Table 1). The response was more pronounced in the more abundant biochemically related Z9 and Z11 isomers compared to the less abundant and biochemically separated Z7-16:OAc (Fig 2). The selection response was characterized by an immediate response in the direction of selection (from generation 0 to generation 1), followed by a gradual progression towards increasing/decreasing relative amounts in the respective selection lines. In the low line, the selection response flattened after cessation of selection, while in the high line, the average log-ratio of Z11:16:OAc to 14:Ald was as high or higher in generations > 10 compared to < 10 (Fig 2D).

**Table 1.** Response of pheromone components in the selected and control lines.

|  | **Low** | **Control** | **High** |
| --- | --- | --- | --- |
| **Total amount** | -20.26 | -50.65** | -22.55 |
| **16:Ald** | -0.02 | -0.11** | -0.05* |
| **Z9-16:Ald** | -0.17*** | -0.15** | -0.08** |
| **Z11-16:Ald** | -0.12*** | -0.14** | -0.08*** |
| **Z7-16:OAc** | -0.19*** | *0.11* | *0.32\*\*\** |
| **Z9-16:OAc** | -0.31*** | -0.02 | *0.2\*\*\** |
| **Z11-16:OAc** | -0.28*** | *0.01* | *0.27\*\*\** |
| **Z9-16:OH** | -0.07* | -0.19** | -0.1** |
| **Z11-16:OH** | -0.12*** | -0.29*** | -0.21*** |

Values represent difference in mean between generation 0 + 1 and generation 9, the last generation following active selection. Components under selection are shown in italics. T-test \* $P < 0.05$ , \*\* $P < 0.01$ , \*\*\* $P < 0.001$ .

**Figure 2.**
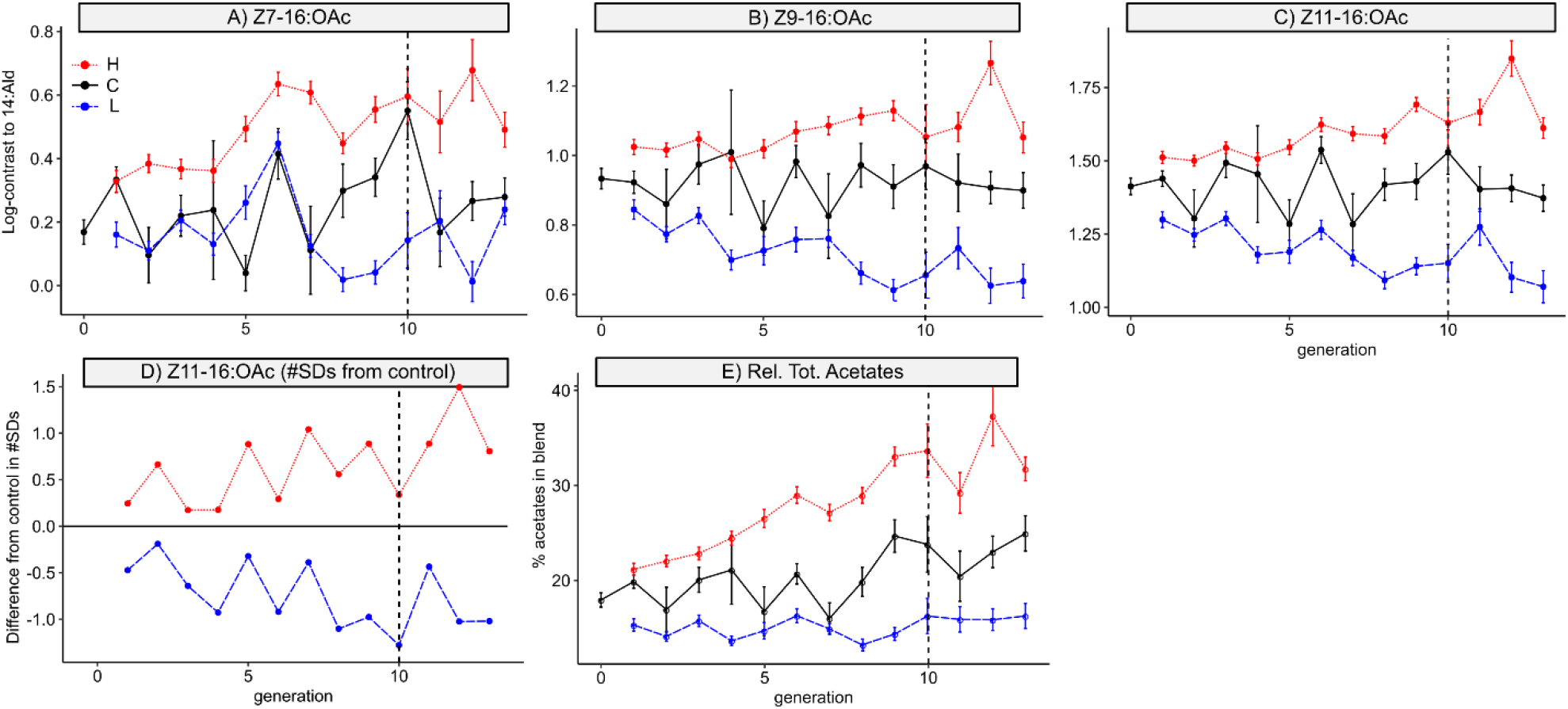
Selection response in the acetates. Values shown are mean ± SEM. The control line is indicated in black, the high line in red (dotted) and the low line in blue (dashed). The dashed vertical line indicates the generation with adult females that developed from the last offspring that went through selection. A – C: Selection responses of the three acetates log-contrasted to 14:Ald. For the control line, 111 females were phenotyped in generation zero, 71 in generation one, and between 6 and 36 for the remaining generations. For the low and high line, respectively 189 – 296 and 189 −295 females per generation were phenotyped. D: Difference in phenotypic means between selection lines and control in units of starting population standard deviation. E: Selection response of the relative total amount of acetates.

The other pheromone components showed a significant decrease over time in both selected and control lines, but no differentiation among selection lines or between selected and control lines (Table 1, Fig 3A-C). The total amount of pheromone measured across the 11 biologically active components remained constant through time in the selected lines, indicating that changing ratios were independent of the pheromone titer (Fig 3D).

**Figure 3.**
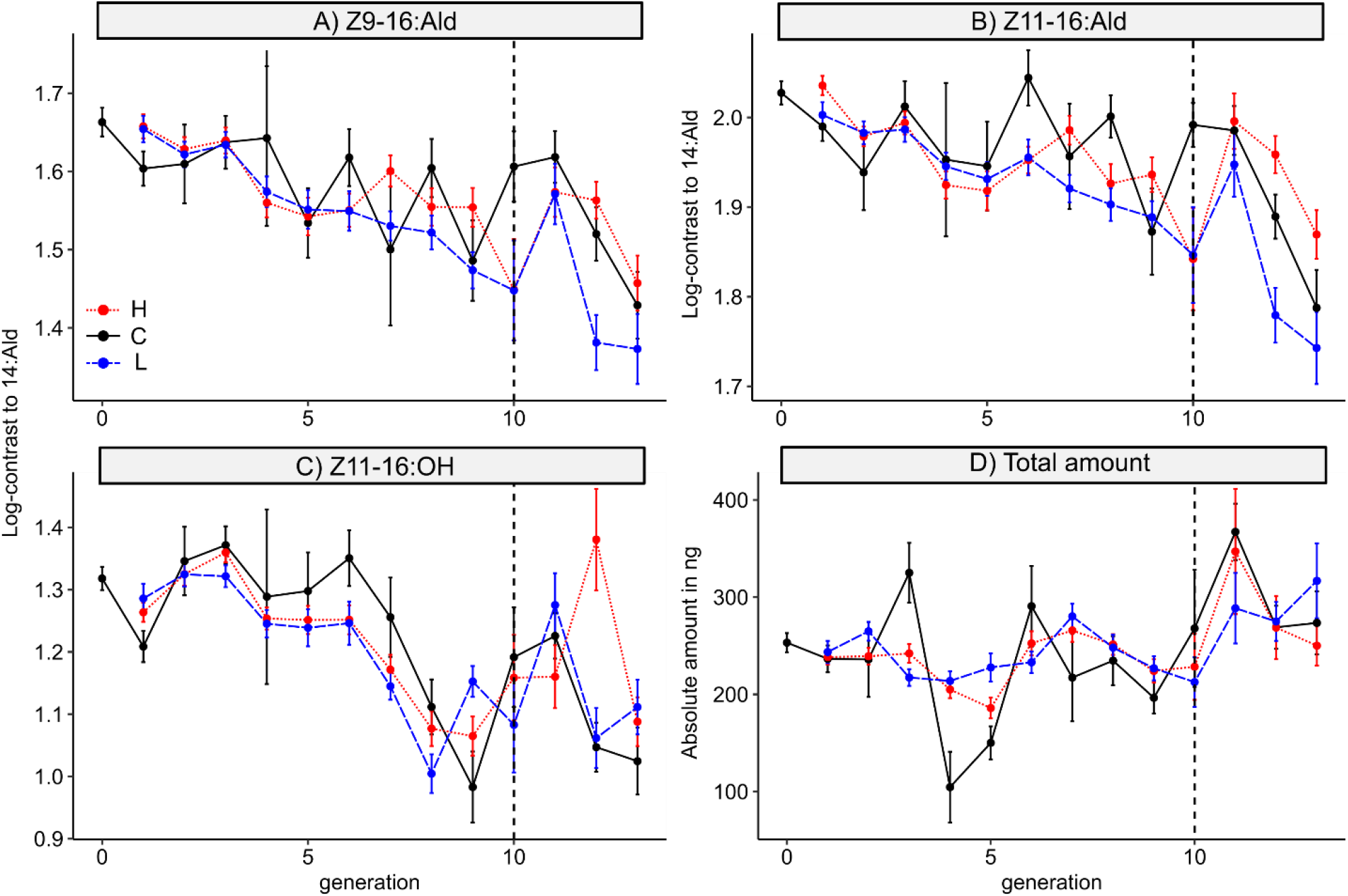
Indirect selection response. Values shown are mean ± SEM for log-contrasts to 14:Ald of the three pheromone components that make up the minimal blend for male attraction and for the absolute amount of pheromone across all 11 biologically active components. The control line is indicated in black, the high line in red and the low line in blue. Dashed vertical lines indicate the generation with adult females that developed from the last offspring that went through truncation selection. Sample sizes as in Fig 2.

### Selection response in genetic (co)variances

In testing whether the selection response was associated with (i) a reduction in genetic variance in Z11-16:OAc (a change in the diagonal elements of the G matrix) and (ii) re-orientation of genetic covariances among pheromone components (changes in the off-diagonal elements of the G matrix), we found no evidence for decreasing genetic variance in Z11-16:OAc or in the other components (Fig A1 in the appendix).

In contrast to genetic variances, we did find changes in the genetic covariance structure across generations. The first two genetic principle components showed a non-significant trend (differences in the mode, but overlapping 90% HPD intervals) towards change in magnitude of their eigenvalue, i.e. the amount of genetic variance they describe (Fig A2). More of the total genetic variance across the four components tended to be in the direction of gPC1 in the high and low lines (posterior mode ranging between 80 and 90%) compared to the starting generation (70%). This means that more genetic variance of the sex pheromone was oriented towards a single dimension in phenotypic space.

Moreover, variation in the component under selection, Z11-16:OAc, was more strongly associated with gPC2 and less with gPC1 in generations after compared to before selection, while the other traits showed a trend in the opposite direction (Fig 4a). Specifically, loadings for Z11-16:OAc on gPC1 were strongly negative prior to the onset of selection, while HPD intervals overlapped zero in both the early and final generations during selection (Fig 4b). Loadings for the Z11-16:OAc on gPC2 were close to zero before, but not after the onset of selection. Loadings for the major component, Z11-16:Ald and its isomer Z9-16:Ald on gPC2 showed the opposite pattern on gPC2 (Fig 4b). These changes in the G matrix resulted in a more modular pheromone blend, because Z11-16:OAc, which was under selection here, became uncoupled from the components that were not under selection.

**Figure 4.**
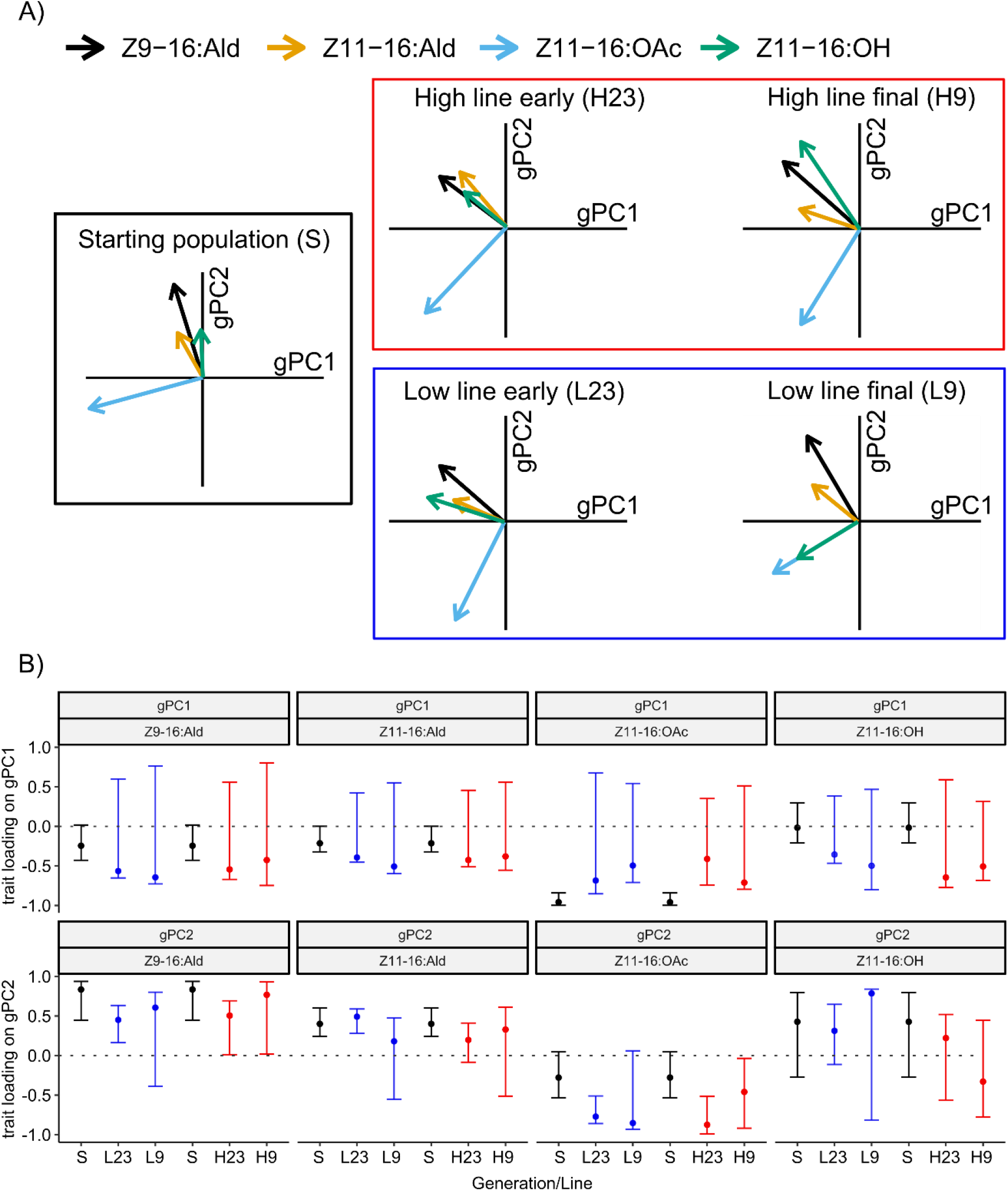
Eigen-analysis of genetic principle components. A) For the starting populations (generations 0 and 1), early generations (2 and 3) and the final generation (9) during active selection, the posterior mode of the correlation between each of the pheromone ratios and the first and second genetic principle components is illustrated by the direction and size of the arrows. B) Trait loadings on genetic principle components. The posterior mode (dots) and 90% HPD interval (error bars) of the loading of each of the log-contrasts on gPC1 gPC2 is shown.

## DISCUSSION

In this study, we investigated how the selection response depends on and shapes the genetic architecture of a multicomponent sex pheromone signal. Through truncation selection, we gradually increased and decreased the relative amount of acetates in the sex pheromone blend of *H. subflexa* across 10 generations. As we hypothesized, we found that the acetates responded readily to selection, diverging three-quarters of a standard deviation away from the control line in 10 generations in both the high and low lines. However, in contrast to our hypothesis that selection would also result in indirect responses in the other traits, we found that the response was limited to the acetates only. In addition, we found that levels of genetic variance did not decrease, opposite to the expected erosion of genetic variance in response to selection. Lastly, in line with our expectations, the genetic covariance structure changed in a way that likely facilitated a selection response in the pheromone. We discuss each of these results further below.

### Univariate responses in multicomponent signals

In nature, sexual signals are often found to be subject to multivariate stabilizing and directional selection (Bentsen et al., 2006; Blankers et al., 2015; Brooks et al., 2005; Devigili et al., 2015; Fisher et al., 2009; Gerhardt and Brooks, 2009; Hine et al., 2011; Oh and Shaw, 2013; Ryan and Rand, 2003; Tanner et al., 2017). Genetic covariances among signal components may pose constraints on the response to selection if the (in)direct selection response in a component conflicts with pressure from other sources of selection on that component. We applied a univariate selection gradient on the relative amount of acetates in the female *H. subflexa* sex pheromone to test whether in the absence of other selection pressures, indirect selection responses are observed in the other sex pheromone components. We justified this univariate selection based on the geographic variation in the relative amount of Z11:16OAc in *H. subflexa*. Females are likely specifically selected for higher rates of acetates to avoid heterospecific mate attraction in regions where the congener *H. virescens* occurs (Groot et al., 2009a). In regions where *H. virescens* is absent, acetate ratios are much lower (Groot et al., 2009a), which suggests there may be costs (life history trade-offs) associated with acetate levels in nature.

We observed that Z11-16:OAc and the other acetates responded readily to selection. Levels of divergence in acetates between the selection lines were similar to or in excess of levels of divergence across the range of *H. subflexa*. After 10 generations in our selection experiment, the high line individuals had on average almost 20% more Z11-16:OAc compared to the low line (Fig 2E), while *H. subflexa* populations in the eastern US were found to have blends with up to 10% more Z11-16:OAc compared to populations in the south-western US and Mexico (Groot et al., 2009a).

The selection response that we found was also surprisingly univariate, as only the three acetates and none of the other components showed divergence between the high and low lines. This indicates that the acetates can evolve independently from the other components. This is surprising, both in the context of the observed genetic correlations in the starting population and in later generations (Fig 4) and in the context of what is known about the shared biochemical pathways across the pheromone components (Groot et al. 2009a; Fig 1). For example, Z11-16:Ald (the major component) and Z11-16:OAc are both biochemically derived from Z11-16:OH (Fig 1; Jurenka 2003) and covaried genetically before and during selection, yet Z11-16:OAc evolved independent from Z11-16:Ald (Fig 2, Fig 3). The independent evolution of acetates and the major component is also biologically important, as the major component is expected to be under stabilizing selection, so directional selection on the acetates in combination with positive covariance between acetates and the major component would result in conflicting selection pressures and constrained selection responses. In contrast, we thus observed independent evolution of coupled traits, which is not an uncommon result of artificial selection experiments (Hill and Caballero, 1992; Saltz et al., 2017), including for sexual signals (Ritchie and Kyriacou, 1996).

### Evolution of the genetic covariance structure

Changes in genetic covariances are rarely documented in natural or laboratory selected populations. Here we found divergence (differences between before and after selection) in the loadings of the pheromone components on the genetic principle components, in particular in the magnitude of the correlation between the component under selection and gPC1 and gPC2. The observed divergence in the genetic covariance structure of the female sex pheromone may result from various mechanisms.

First, the covariance structure may change due to changes in genetic variances, because stabilizing or directional selection is expected to erode genetic variance (Barton and Turelli, 1989; Estes and Arnold, 2007). Although our truncation selection increased and decreased the phenotypic mean by three-quarters of a standard deviation in the high and low line, respectively, we observed no significant decrease in the coefficient of additive genetic variance. One explanation is that several genetic factors may underlie the trait under selection and that selection therefore leaves no detectable signature on the genetic variance associated with that trait (Bulmer, 1971; Johnson and Barton, 2005). Genetic mapping studies have revealed several QTL underlying difference in sex pheromone composition in *H. subflexa* and even a single conversion step in the biochemical pathway, such as from Z11-16:OH to Z11-16:OAc, could be catalyzed by multiple enzymes (Groot et al., 2013). In addition, variation in pheromone composition in *H. subflexa* is explained by fitness variation (Blankers et al., 2021), indicating many different genetic factors likely contribute small additional fractions to the total variance. Another explanation comes from population genetics: re-orientation of genetic covariances in response to bottlenecks may free up additive genetic variance, thereby paradoxically increasing levels of genetic variance (Carson, 1990; Templeton, 2008). Since truncation selection is effectively a non-random bottleneck of the population, this may also explain the maintenance of genetic variance that we observed. Lastly, to avoid inbreeding depression, we maintained large populations and avoided mating first and second-degree relatives. Such a mating scheme likely counteracts some loss in genetic variability due to selection (Du et al., 2021).

Second, the genetic covariance structure can evolve due to changes to interactions among unlinked genetic loci affecting the co-expression of multiple traits (Wolf et al., 2005). Since there is a variety of enzymes involved in the conversion between pheromone components (Groot et al., 2016; Jurenka, 2003; Tillman et al., 1999) and variation in different pheromone components is due to effects at different chromosomes (Groot et al., 2009b; Groot et al., 2014), it is likely that a significant portion of genetic covariance among sex pheromone components results from interactions among unlinked loci. It may therefore be unsurprising that we see an immediate change in the covariance structure following the onset of selection. This finding is of broad significance to predicting selection response from quantitative genetics, because these predictions often depend on assumptions of a constant matrix over multiple generations. Our findings lend support to both theoretical (Arnold et al., 2008; Eroukhmanoff, 2009; Roff, 2000) and empirical (Björklund et al., 2013; Hine et al., 2009; Uesugi et al., 2017) research on the potential instability of the G-matrix.

Interestingly, we found a relationship between phenotypic evolution and change in the genetic covariance structure. The component under selection, Z11-16:OAc was decoupled from the other components as observed by divergence in the loadings on gPC1 and gPC2. This observation thus fits the prediction that genetic variances and covariances may be reoriented to align the phenotype with the dominant direction of selection (Melo and Marroig, 2014). These changes in the G matrix can likely facilitate a more specific response where only Z11-16:OAc evolves in the absence of indirect selection responses. The observed evolution of the G matrix is biologically relevant, because selection for heterospecific mating avoidance favors higher rates of acetates in the female sex pheromone of *H. subflexa* (Groot et al., 2006) while intra-specific (sexual) selection is expected to act stabilizing on the rates of the other major and critical secondary sex pheromone components.

### Conclusion

From our results, we conclude that: 1) pheromone components in *H. subflexa* can evolve in response to selection independent of other components of the sex pheromone; 2) selection need not erode the genetic variance in order to drive this phenotypic change; 3) selection alters the genetic correlations among pheromone components. Our study thus shows that univariate selection responses in multicomponent sexual signals are possible, even though genetic correlations were high both prior to and during selection. Our study also shows that sexual signal components can respond to selection without reductions in genetic variance, but with rapid changes in the genetic covariance structure. These results correspond to geographic variation in the sexual signal found in this species and help to explain how sexual signals in general can respond to selection without running into constraints from indirect selection responses or depleted genetic variation. This may be one reason why sexual signals have evolved rapidly and repeatedly throughout the evolutionary history of animal taxa, thereby contributing to the speciation process.

## ACKNOWLEDGEMENTS

We thank Dennis van Veldhuizen for his invaluable contribution to the rearing of our moths. We also thank Laura Caton, Frederica Lotito, and Lilian Seip for their help with the selection lines. We further thank Jet ten Berge for digitizing the pedigree, Michiel van Wijk for commenting on a previous version of the manuscript, and all members of the Groot lab for their helpful and insightful discussions.

## AUTHOR CONTRIBUTIONS

All authors contributed to the conceptualization of the study; TB, EF, and EBS performed the experiments; TB analyzed the data; and all authors contributed to writing the manuscript.

## DATA ACCESSIBILITY

Data and analysis scripts will be archived in Dryad.

## APPENDIX

**Table A1.**
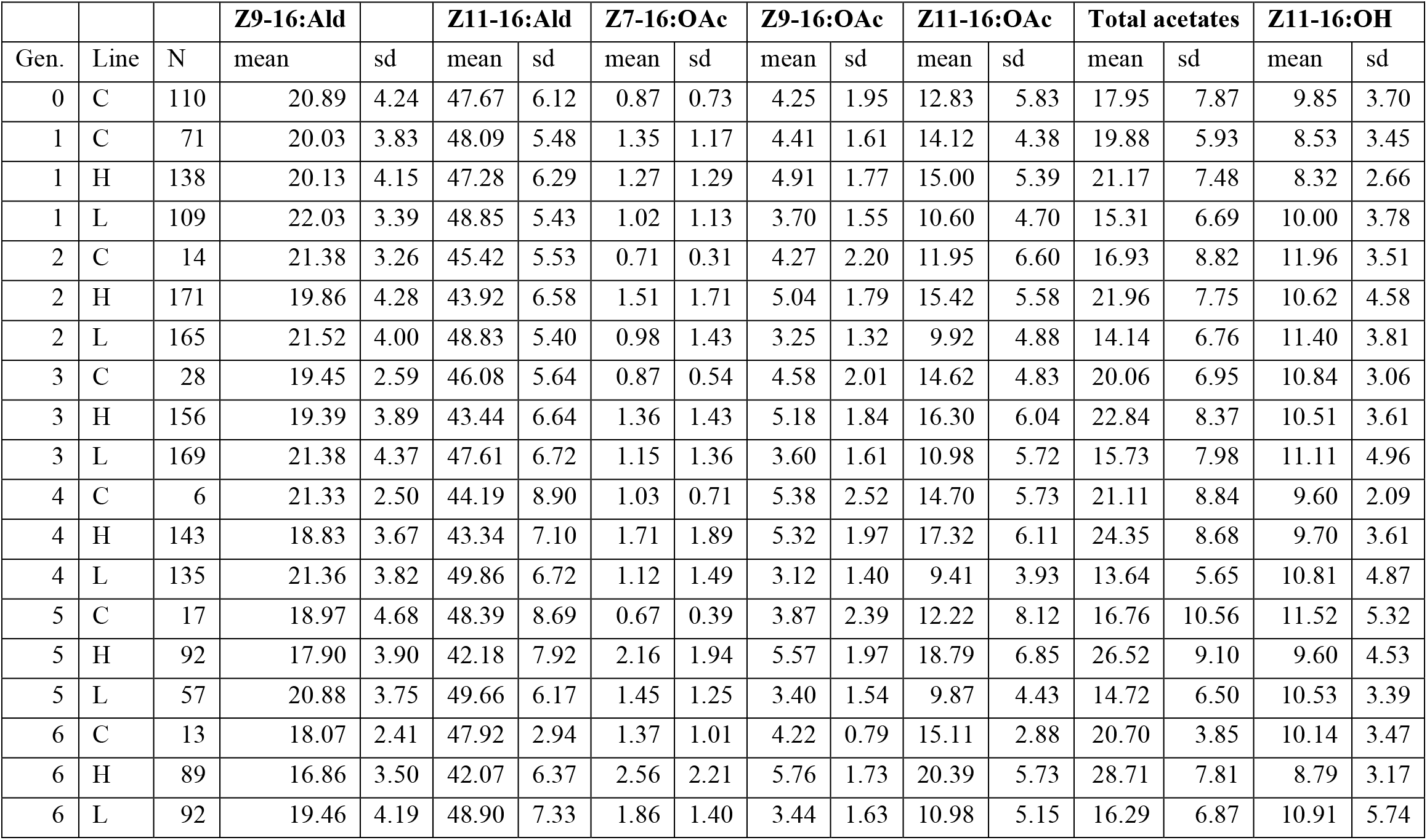
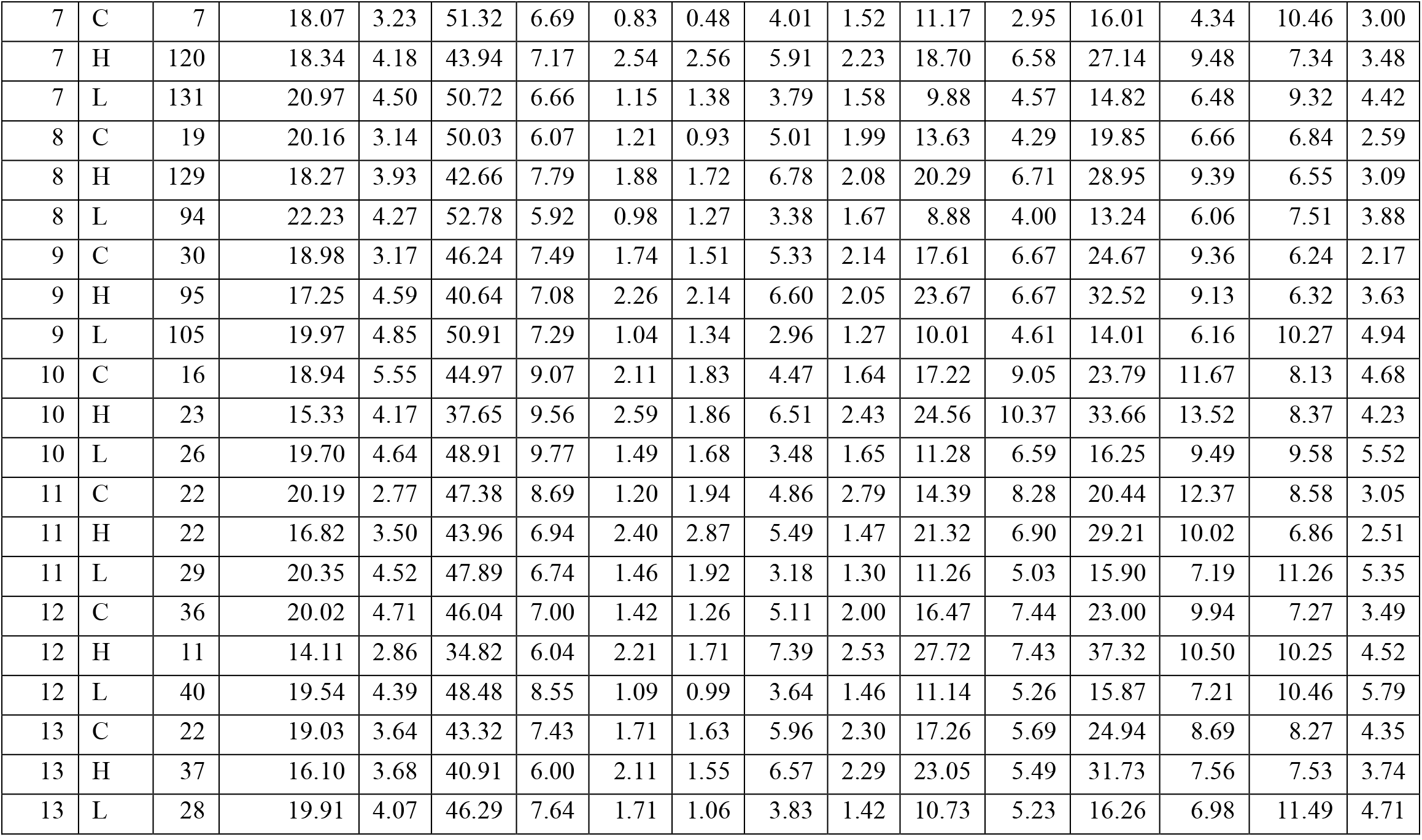
Relative amounts for the analyzed components as well as all three acetates and their sum along with sample sizes for each line and each generation.

**Figure A1.**
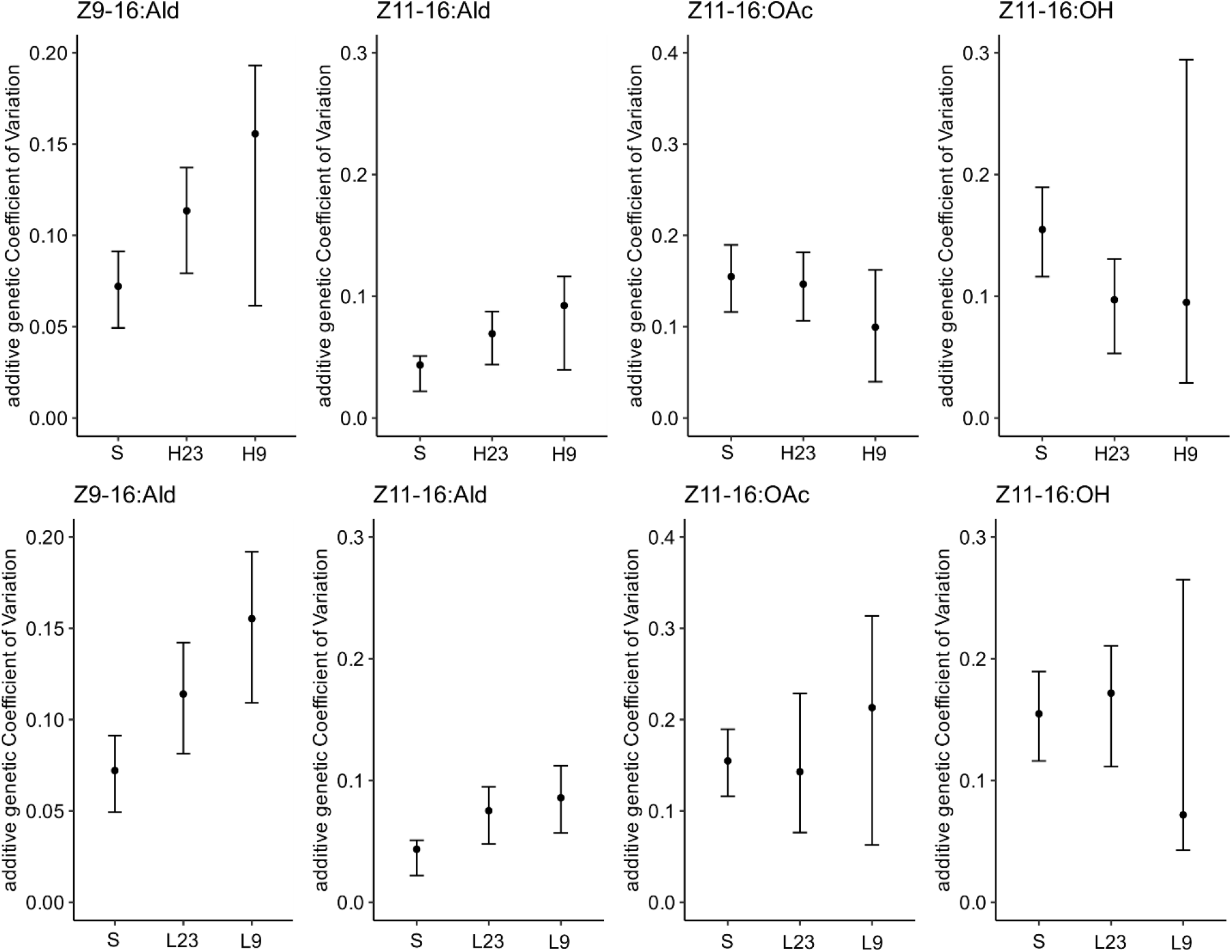
Additive genetic coefficients of variation for starting populations, early generations and final generation. The posterior mode (dots) and 90% HPD interval (error bars) of the coefficients of variation are shown.

**Figure A2.**
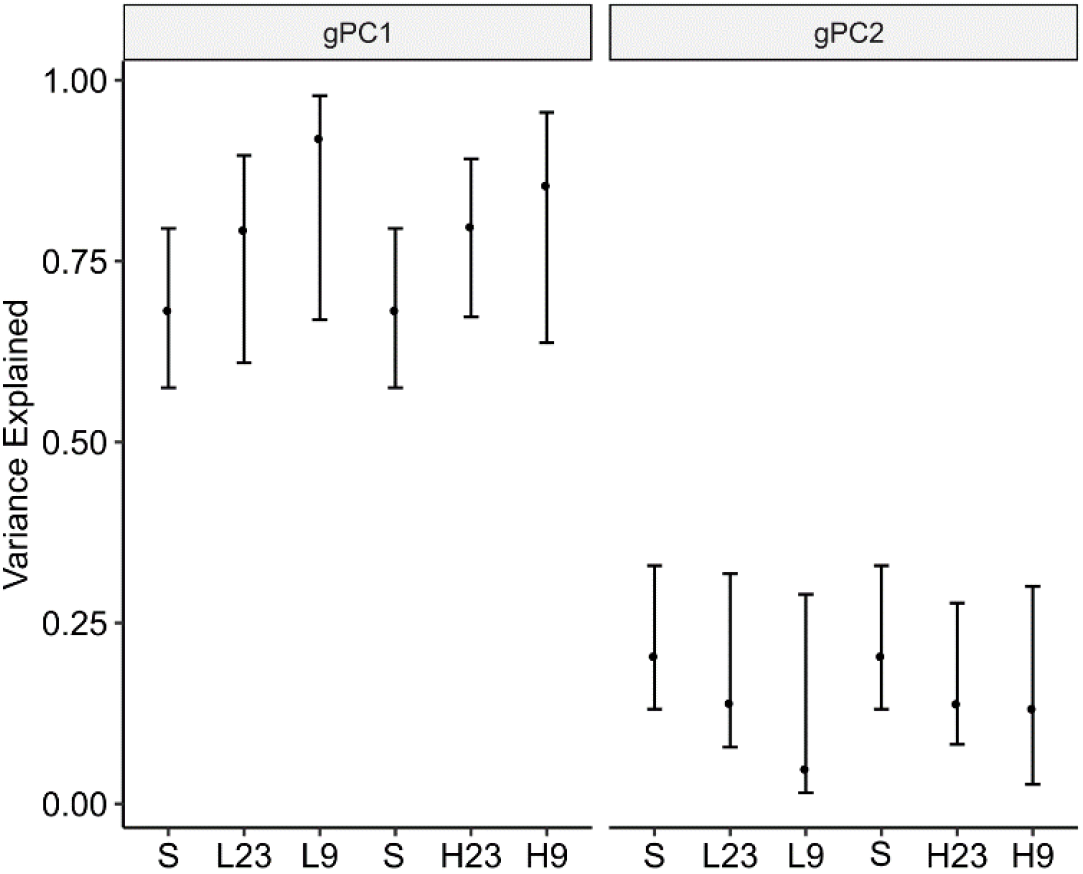
Variance explained by gPC1 and gPC2 for the starting populations and the early and final generations during selection in the High and Low lines. The posterior modes (dots) and 90% HPD intervals (error bars) of the variance estimates are shown.

